# Dark-based Optical Sectioning assists Background Removal in Fluorescence Microscopy

**DOI:** 10.1101/2024.03.02.578598

**Authors:** Ruijie Cao, Yaning Li, Wenyi Wang, Guoxun Zhang, Gang Wang, Yu Sun, Wei Ren, Jing Sun, Yiwei Hou, Xinzhu Xu, Jiakui Hu, Yanye Lu, Changhui Li, Jiamin Wu, Meiqi Li, Junle Qu, Peng Xi

**Author notes:** Correspondence should be addressed to J. Q. and P.X. These authors contribute equally to the work (Ruijie Cao, Yaning Li).

## Abstract

A fundamental challenge in fluorescence microscopy is the defocused background caused by scattering light, optical aberration, or limited axial resolution. Severe defocus backgrounds will submerge the in-focus information and cause artifacts in the following processing. Here, we leverage a priori knowledge about dark channels of biological structures and dual frequency separation to develop a single-frame defocus removal algorithm. It stably improves the signal-to-background ratio and structural similarity index measure of images by approximately 10-fold, and recovers in-focus signal with 85% accuracy, even when the defocus background is 50 times larger than in-focus information. Our Dark-based optical sectioning approach (Dark sectioning) is fully compatible with various microscopy techniques, such as wide-filed microscopy, polarized microscopy, laser-scanning / spinning-disk confocal microscopy, stimulated emission depletion microscopy, lightsheet microscopy, and light-field microscopy. It also complements reconstruction or processing algorithms such as deconvolution, structure illumination microscopy, and super-resolution optical fluctuation imaging.

## Introduction

Fluorescence microscopy plays an important role in cell biology, as the fluorescently-labeled signals can specifically represent target organelles to visualize the subcellular structure and intercellular interactions^1-4^. However, defocused backgrounds caused by scattered light, optical aberrations, or defocused axial fluorescence^5-7^ can severely contaminate the in-focus signal and cause artifacts in subsequent algorithmic processing in fluorescence microscopy such as wide-field (WF) microscopy^8^. Although optical approaches such as laser-scanning confocal microscopy^9^ or multi-photon microscopy^10^ provide enhanced optical sectioning performance, they have relatively lower temporal resolution with more photobleaching due to high doses of light. Other imaging techniques relying on the active illumination modulation and WF-based array detection, such as spinning-disk confocal, multi-focal structured illumination microscopy, three-dimensional structured illumination microscopy (SIM)^11, 12^, lightsheet microscopy^13^, or super-resolution optical fluctuation imaging^14, 15^ (SOFI), also suffers from the defocused noise and lower imaging speed despite improving axial sectioning.

Computational approaches have become increasingly important for improving the performance of optical microscopy. Typical computational fluorescence approaches include reconstruction algorithms such as high-and-low-frequency microscopy (HILO), SIM^16, 17^, SOFI^14^, and pre/post-processing algorithms such as deconvolution^18, 19^ and denoise^20^. Nearly all these methods require pre-background removal to avoid severe artifacts in the final image^5^. However, their background removal parts are mostly based on the assumption that out-of-focus information lies in the low-frequency while in-focus information lies in the high frequency, just like rolling ball^21^ or sliding paraboloid^22^ algorithms. These approaches have issues with incorrect or incomplete defocus background removal, as well as low weak signal retention. Advanced open-source background removal algortithms such as deepWonder^23^, BF-SIM^5^, QLFM^24^ specific to SIM, widefield (for calcium imaging only), or light-field, respectively. Popular deep learning methods for background removal can perform the task, but they lack transparent processes, and the performance is heavily relying on the training dataset and parameter tunning^25^. Given the trade-offs between physical sectioning methods and the growing need for computational fluorescence image processing, exploring the universal features of fluorescence background and proposing a powerful background removal method is urgently required for fluorescence microscopy.

Inspired by natural image de-haze processing, we leverage the dark channel priority for out-of-focus fluorescent background removal. Dark channel priority is one of the most successful methods in natural outdoor image dehazing^26, 27^. This simple yet powerful principle significantly advanced single-image dehazing. However, it requires even background distribution and color characteristics of outdoor scenes. Complex defocus structure and uneven background distribution in fluorescence images, where the pixel grayscale represents the fluorescence emission intensity, preclude dark channel prior in direct application which would sacrifice fine details and incompletely remove local backgrounds. While Dark channel priority can distinguish and eliminate haze in outdoor images, it fails in the application on fluorescence.

Here we present Dark sectioning, a versatile algorithm that effectively segments and removes out-of-focus fluorescence background by exploiting physical characteristics, dark channel prior and dual frequency separation. Dark sectioning eliminates defocused background while preserving weak in-focus signals with high fidelity and stability for major fluorescence techniques including widefield, confocal, lightsheet, HiLo, 2D/3D/polarized-SIM, STED and SOFI microscopies. It can elevate wide-field microscopy results to reach confocal optical-sectioning capability, improve one-photon to two-photon results, and 2DSIM to 3DSIM results. Dark Sectioning is fully compatible to various microscope setups and reconstruction/processing algorithms. Thus, it provides a widely applicable solution to markedly improve axial sectioning in fluorescence microscopy.

## Results

### Principle and performance of Dark sectioning

Dark channel priority was first proposed for haze removal in natural images. The core of dark channel priority is that most local blocks in haze-free outdoor images contain some pixels with very low intensities in at least one RGB channel, and the dark channel *dark* [*I*(*x, y*)] of the image *I* (*x, y*) can be expressed as^26^:

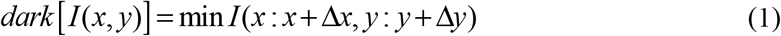

Where (Δ*x*, Δ*y*) denotes the block size, and the dark channel can be expressed as the minimum value of RGB in a certain block size.

Different from traditional Dark-based haze-removal methods, the dark channel can also distinguish in-focus and out-of-focus information when the in-focus object has a single color. As shown in Fig. 1(a), an almost zero dark channel is presented in the in-focus yellow flower, but the dark channel of the out-of-focus leaves exhibit blocky fluctuating non-zero values. Then, we try to conduct the dark channel operation of WF images and find that the dark channel of the out-of-focus part is almost equal to the background distribution in Fig. 1(b). This is because that out-of-focus background can be expressed as the convolution between the ground-truth (GT) image and the out-of-focus point spread function (PSF), and the out-of-focus PSF is larger than the in-focus PSF due to the physical characteristics of microscopy. So, the dark channel of in-focus PSF exhibits all-zero values while the dark channel of out-of-focus PSF exhibits none-zero in Fig. 1(c). We define this method to distinguish in-focus and out-of-focus information as the core of dark priority in fluorescence images, and Dark sectioning is developed. Hence, we redefine the block size in the dark channel as:

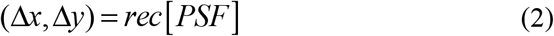

**Figure 1.**
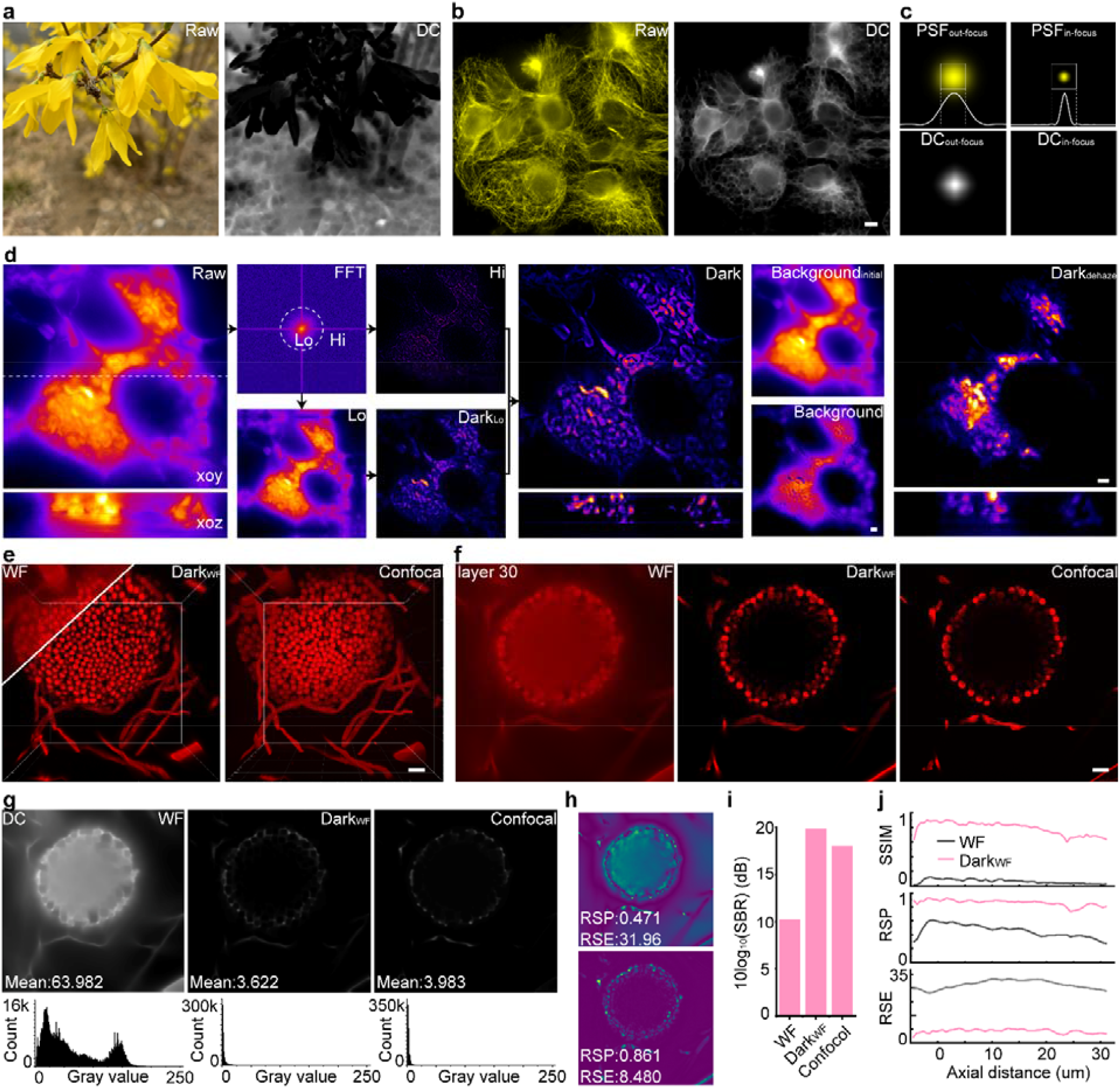
Principle and performance of Dark sectioning. (**a**) The dark channel of natural image with big aperture, including the in-focus flower and out-of-focus leaves. (**b**) The WF image of tubulin structure in U2OS cells with strong out-of-focus background and its corresponding dark channel image. (**c**) Comparisons of in-focus PSF and out-of-focus PSF with their corresponding dark channel. (**d**) The simplified algorithm flow of Dark sectioning using a WF image of the mouse tail section as an example, including the separation of high-frequency and low-frequency parts, the Dark section processing of the low-frequency part, final results, estimated background, and the extracted background. Comparison between WF, Dark_WF_, and Confocal of aspergillus conidiophores in (**e**) multi-layer view and (**f**) single-layer view in 30^th^ layer, with (**g**) their corresponding dark channel images (upper) and histograms (bottom). (**h**) The RSP and RSE of WF and Dark_WF_ in (**f**) using Confocal as GT. (**i**) The comparisons of SBR between WF, Dark_WF_, and Confocal images. (j) The SSIM, RSP, and RSE between WF, Dark_WF_, and Confocal images along the axial distance. Lateral scale bar: 4 μm. Axial scale bar: (**d**) 209 layers, 125 nm per layer, (**f**) 60 layers, 0.7 μm per layer.

Where *rec*[·] denotes the size of the external rectangle which is slightly larger than PSF, which can better distinguish in-focus and out-of-focus information with the physical foundation.

As is known to all, the background mainly exists in the low-frequency part of the image. So, we divide the image into the low-frequency part *I*_*Lo*_ (*x, y*) and the high-frequency part *I*_*Hi*_ (*x, y*), and only use the low-frequency part for Dark sectioning to better preserve the weak information as shown in Fig. 1(d). Then, Dark sectioning further divides the image into the background part *A*(*x, y*) and the no-background part *J* (*x, y*) with a certain transmission ratio *t* (*x, y*) :

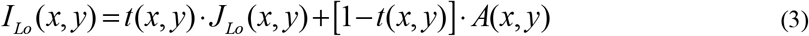

Furthermore, we introduce an uneven background rather than a fixed atmosphere in dehazing algorithm to better fit the background distribution of fluorescence images, quantify global defocus information by adjusting parameters, and use iterative removal for local and global background. Because of the assumption that the high-frequency part of the background removed image is the same as the raw image, namely *J*_*Hi*_ (*x, y*) *I*_*Hi*_ (*x, y*) in Fig. 1(d). Hence, the final image *J* (*x, y*) with the background removed can be obtained:

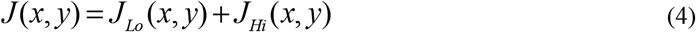

Where the initial background *A*(*x, y*) and the raw image minus the background removed image *I* (*x, y*) −*J* (*x, y*) (defined as background). After Dark sectioning, the removed background is more detailed, so the final result preserves the details of the sample as shown in Fig. 1(d). And the traditional dehazing algorithm based on dark channel priority (Dark_dehaze_) to remove the background of the mouse tail section is shown in the left part of Fig. 1(d). Through comparison between Dark sectioning and Dark_dehaze_ illustrates that Dark_dehaze_ does not apply to fluorescence images because of the loss of weak information and incorrect removal of background.

Because confocal microscopy^28^ with a point scanning imaging method has an improved resolution in the z-axis and greatly reduced defocused background, we further did the comparison between WF, Confocal, and WF images processed by Dark sectioning (Dark_WF_) of aspergillus conidiophores as shown in the multi-layer view in Fig. 1(e) and single-layer view (the 30^th^ layer) in Fig. 1(f). Optical sectioning capability of Dark sectioning is shown and validated by the confocal image, and the corresponding dark channel images are shown in Fig. 1(g). It can be seen that Dark_WF_ and confocal images with optical sectioning capability can concentrate histogram of dark channels close to 0 with the mean value of dark channels being 3.622 and 3.983 in an 8-bit image, far less than 63.982 in WF. This proves the dark channel priority and the correctness of the removed background.

We use the NanoJ plugin^29^ to evaluate the fidelity of WF and Dark_WF_ compared with confocal images in Fig. 1(h), finding that Dark sectioning can improve the resolution-scaled Pearson coefficient (RSP) from 0.471 to 0.861 and reduce resolution-scaled error (RSE) from 31.96 to 8.48, and improve the signal-to-background (10log_10_(SBR)) of WF from 10dB to 20dB, namely 10 times, in Fig. 1(i). And, in Fig. 1(j), we further compare the structural similarity index measure (SSIM), RSP, and RSE between WF and Dark_WF_ in different axial layers using confocal image as GT, finding the SSIM has been stably improved by about 10 times with the maximum SSIM with confocal images exceeds 0.9. And the RSP and RSE have been improved by 2 times and 4 times correspondingly. This demonstrates that Dark sectioning can achieve a high-fidelity optical sectioning effect similar to confocal images only based on WF images.

### High-fidelity of Dark sectioning

To further validate the fidelity of Dark sectioning, we first do the simulation using a hollow sphere and linear structure in Extended Data Fig. 4 and Supplementary Note 5, and we artificially change the proportion of the PSF on different z-axis layers to control the severity of defocused background. Even though the in-focus intensity is 2% of out-of-focus information, Dark sectioning can recover in-focus information to a great extent. Furthermore, we build DMD-based multi-modality joint SIM microscope containing HiLo, 2DSIM^4, 16^, and 3DSIM using a digital micromirror device (DMD) and electro-optics modulator (EOM)^30^ in Extended Data Fig. 5 for cross-validation. HiLo microscopy^31, 32^ has a better optical section capability than WF, but it needs two images with illuminated patterns to reconstruct one image with the background removed. Furthermore, 3DSIM^11, 12^ with double resolution in the *xyz* directions has a great background removal capability. Thus, we do the comparison between WF, Dark_WF_, HiLo, 2DSIM, 3DSIM, and 2DSIM processed by Dark sectioning (Dark_2DSIM_) using mouse kidney section as an example in Fig. 2(a, b). We find that Dark sectioning can correctly remove the defocus background of WF and 2DSIM, which is proved by HiLo and 3DSIM. We further do the NanoJ plugin using 3DSIM as GT to evaluate the fidelity of Dark sectioning. It can be found that Dark_WF_ and Dark_2DSIM_ outperform WF, 2DSIM, and even HiLo images with a higher RSP and RSE in Fig. 2(c). And it can be seen in the fourier domain of 2DSIM and Dark_2DSIM_ that Dark sectioning can greatly suppress the frequency peak caused by defocused background along the white profile in Fig. 2(d), so as to reduce the hexagonal artifacts.

**Figure 2.**
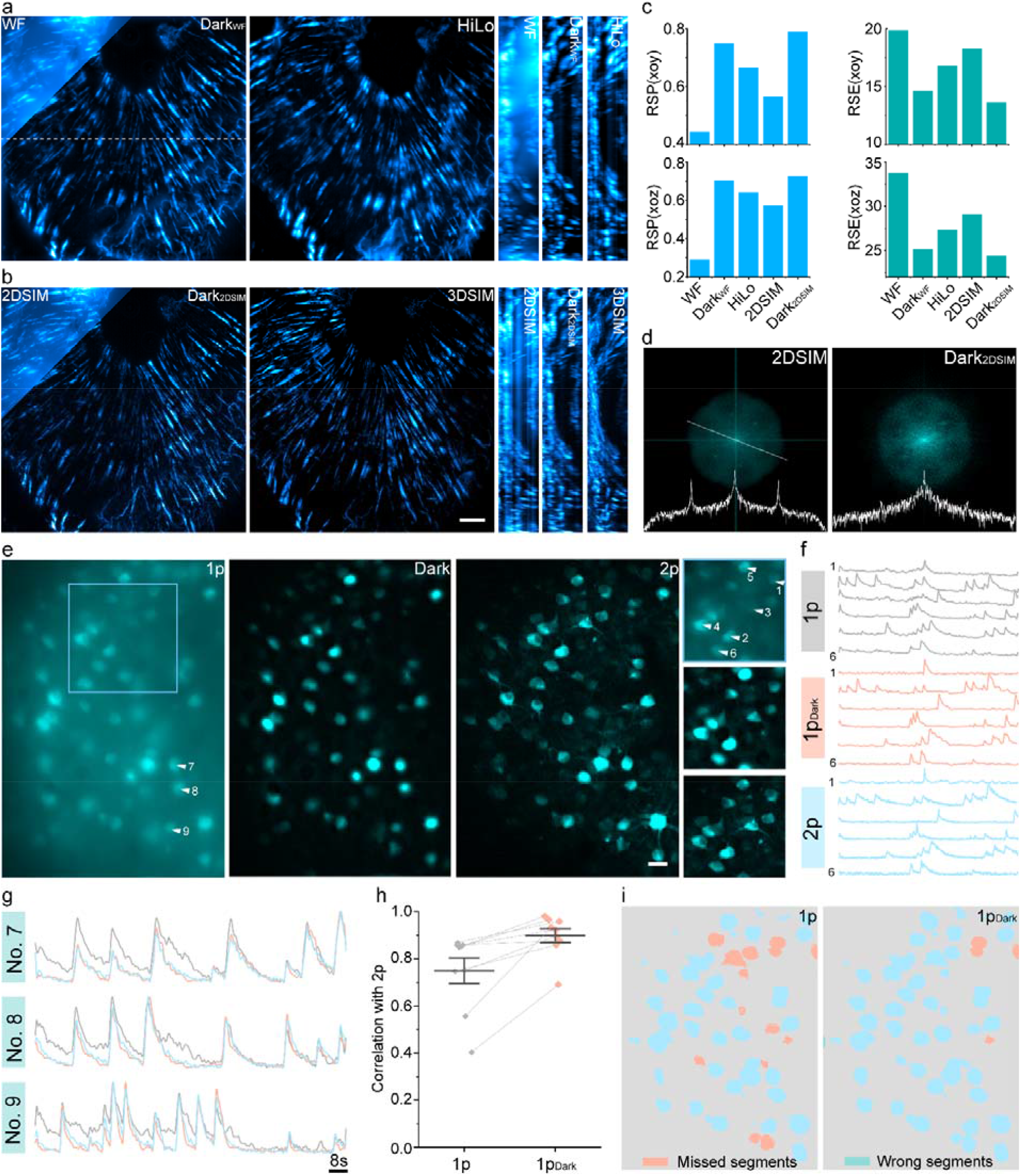
Multi-mode structure illumination microscopy methods and one/two-photon joint imaging modality prove the high fidelity of Dark sectioning. Comparisons between max intensity projection (MIP) images of (**a**) WF, Dark_WF_, and HiLo, (**b**) 2DSIM, Dark_2DSIM_, and 3DSIM of actin filaments in mouse kidney section in *xoy* and *xoz* plane. (**c**) NanoJ analysis of RSP and RSE in respect of WF, Dark_WF_, HiLo, 2DSIM, and Dark_2DSIM_ using 3DSIM as GT in *xoy* and *xoz* plane. (**d**) Comparison between the Fourier domain of 2DSIM and Dark_2DSIM_ with the corresponding intensity profile along the white line. (**e**) MIP images comparison between 1p, 1p_Dark_, and 2p for neurons of mouse brain tissue with their corresponding magnified zones. (**f**) The corresponding temporal neuroelectric activities imaged at 15 Hz per frame (totally 2000 frames) are marked by the triangle for neurons No. 1-6 in (**a**). Gray, pink, and blue lines represent 1p, 1p_Dark_, and 2p images, respectively. (**g**) Zoon-in plots of temporal neuroelectric activities of neurons No. 7-9. (**h**) Temporal pearson correlation coefficients of 1p images using 2p images as GT before (0.749 ± 0.052, mean ± s.d.) and after (0.898 ± 0.023) Dark sectioning based on 1p images. (**i**) Weka segmentation results of 1p and 1p_Dark_ with the corresponding missed and wrong segments. Lateral scaler bar: (**a**) 4 μm, (**e**) 20 μm. Axial scale: (**a**) 39 layers, 125 nm per layer.

What’s more, we also use a custom-built one-photon (1p) and two-photon (2p) joint imaging system^23^ to validate the fidelity of Dark sectioning and the capability to improve the accuracy of neuronal extraction in mouse brain tissue. 1p images are processed by Dark sectioning and 2p images are used as GT. We can find that Dark sectioning can restore the in-focus information in both spatial (Fig. 2(e)) and temporal dimensions (Fig. 2(f)) based on background-overwhelmed 1p images. The better correspondence of temporal neuronal activities between 2p and Dark sectioning processed one-photon (1p_Dark_) images with the selected neuroelectric activities in Fig. 2(g) and the corresponding coefficient in Fig. 2(h) from (0.749 ± 0.052, mean ± s.d.) to (0.898 ± 0.023) illustrates the fidelity of Dark sectioning. Then, the trainable Weka segmentation^33^ of neurons is conducted for 1p and 1p_Dark_ images in Fig. 2(i), and we can find that after Dark sectioning, more neurons submerged by defocus background can be distinguished. Those favorable results show the fidelity and capability to extract neurons of Dark sectioning.

### Dark sectioning for multiple modalities of fluorescence microscopes

Then, we use Dark sectioning into multiple modalities of fluorescence microscopes to prove its wide applications. We firstly introduce it into polarized microscopy and do the comparison between polarized wide-field (pWF), polarized 3DSIM (p3DSIM), polarized WF images assisted by Dark sectioning (pDark_WF_), and polarized 3DSIM images assisted by Dark sectioning (pDark_3DSIM_) of actin filament in mouse kidney section. Dipole orientation interpretation^11, 34^ depends on the intensity ratio of WF images from different polarization angles, and defocused backgrounds change the intensity ratio of WF images, making it difficult to analyze the dipole orientation in Fig. 3(a). Dark sectioning can greatly preserve the polarization information with enhanced polarization precision from 6° to 1.7° as shown in Fig. 3(b) along the white arrow in the magnified zone in Fig. 3(a). The increased precision (pink and yellow arrows) and decreased background make pDark_WF_ acceptable for use with the same polarization precision and lower photobleaching compared with pDark_3DSIM_. Then, we use Dark sectioning for STED microscopy to observe mitochondrial membranes in Fig. 3(c). We can find that the Dark sectioning can help STED images to remove the background and make the cristae clearer as the white arrow shows, which also demonstrates that noise will not affect the performance of Dark sectioning. We also tested Dark sectioning for small-animal imaging (SI) of live mouse vascular and found that Dark sectioning processed small-animal imaging images (Dark_SI_) can effectively remove the scattering background caused by mouse skin or other tissue, greatly restoring the location and change of mouse blood vessels in Fig. 3(d). We also use lightsheet data of *D. melanogaster* embryo^35^ to demonstrate that Dark sectioning can greatly delimitate the background caused by degraded quality in deeper tissue in Fig. 3(e). What’s more, lightsheet data of C. elegans embryos (beam waist being 1.6 μm and 3.3 μm) in Fig. 3(f) shows that Dark sectioning can help lightsheet data with 3.3 μm beam waist to exceed the optical sectioning capability of that with 1.6 μm beam waist. We further use the Bio-LFSR^36^ dataset of light-field microscopy to reconstruct the live mouse liver by virtual-scanning light-field microscopy (Vs-LFM) with or without Dark sectioning. We find that even if Vs-LFM has reduced the motion artifacts of neutrophils with high-speed migration in mouse liver to a great extent, there are still motion artifacts because of the motion of the background. If we use Dark sectioning before deconvolution, the background can be greatly reduced so the final image will have almost no motion artifacts in Fig. 3(g). Benefitting from the reduced motion artifacts, Dark sectioning can help Vs-FLM better observe the 3D dynamic process of the mouse liver with almost no artifacts as shown in Fig. 3(h).

**Figure 3.**
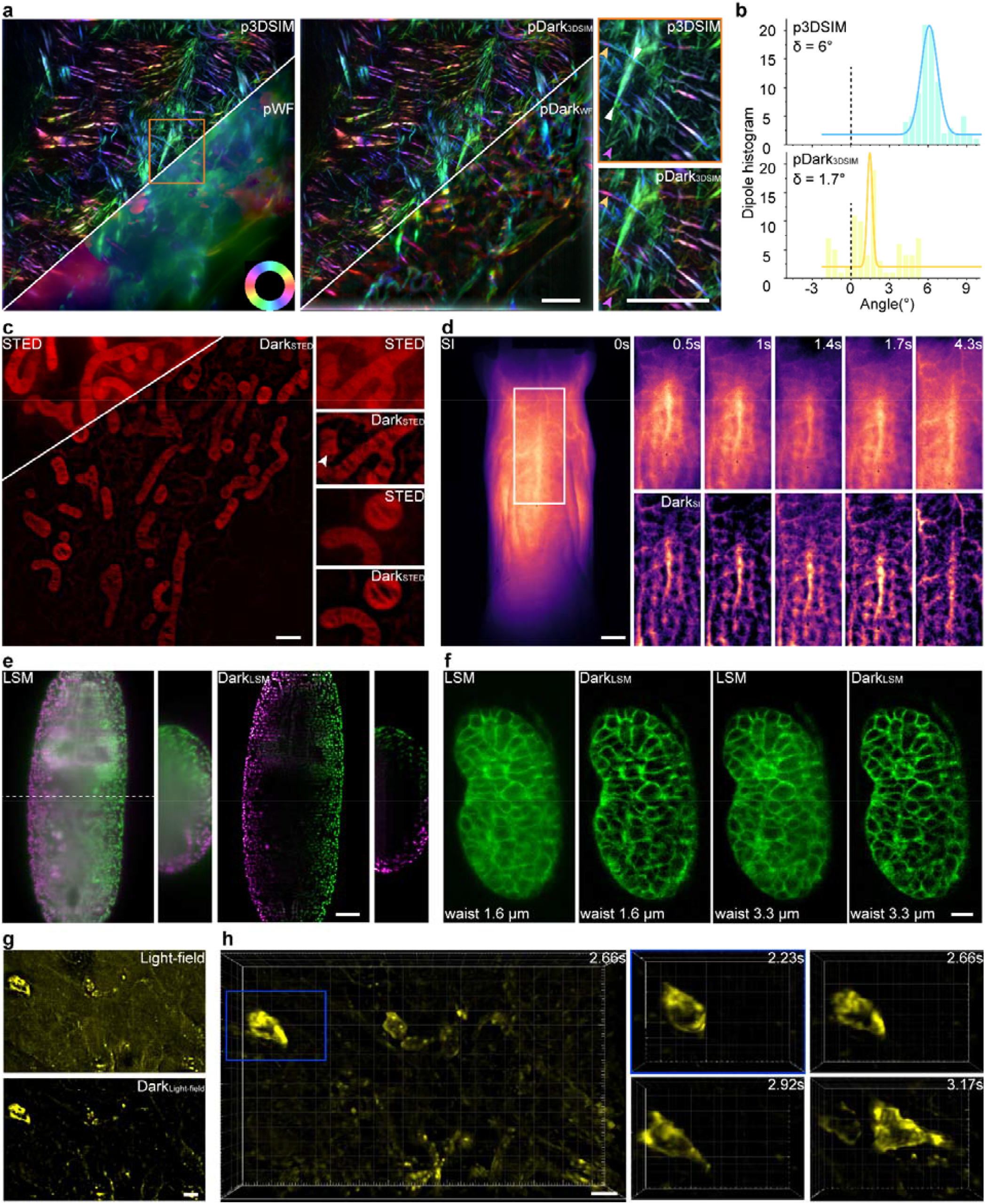
Dark sectioning assists multiple modalities of fluorescence microscopy. (**a**) Dark sectioning helps pWF and p3DSIM microscopy achieve better precision in dipole orientation with the comparisons between p3DSIM, pWF, pDark_3DSIM,_ and pDark_WF_. The yellow and pink arrows in the magnified zone prove the higher precision acquired by Dark sectioning. (**b**) The histogram of deviation between actual and interpreted orientation along the white arrows in (**a**) with the corresponding Gaussian fitting. (**c**) Comparison be STED and Dark_SETD_ to observe mitochondria in Cos-7 cells labelled by SiRPFA after drug-induced apoptosis. (**d**) Small-animal imaging for observing vascular in live mouse with the comparison between SI and Dark_SI_. (**e**) The lightsheet (LSM) and Dark sectioning processed image (Dark_LSM_) of *D. melanogaster* embryo with His2Av-EGFP labeled nuclei. The pink color is the direction of lightsheet from left to right, and the same frame (in green color) was imaged with sequential excitation incident on the embryo from right to left. Also shown is an overlay of the two images with opposing light sheet propagation directions. (**f**) The lightsheet image of *C. elegans* embryos with cell membrane labelled with the comparison of Dark_LSM_, corresponding to the illumination beams (beam waist being 1.6 μm and 3.3 μm). Note that 3.3 μm beams means more out-of-focus fluorescence. (**g**) The light-field image of neutrophils with high-speed migration in a living mouse liver in Bio-LFSR dataset using VsLFM reconstruction with/without Dark sectioning with (**h**) its 3D dynamic process. Lateral scale bar: (**a, c**) 4 μm, (**e**) 0.5 μm, (**f**) 40 μm, (**d**) 2 mm, (**e**) 40 μm, (**f**) 5 μm, (**g, h**) 100 μm. Axial scale: (**e**) 201 layers, 0.48 μm per layer, (**g, h**) 230 nm per layer, 31 layers.

**Figure 4.**
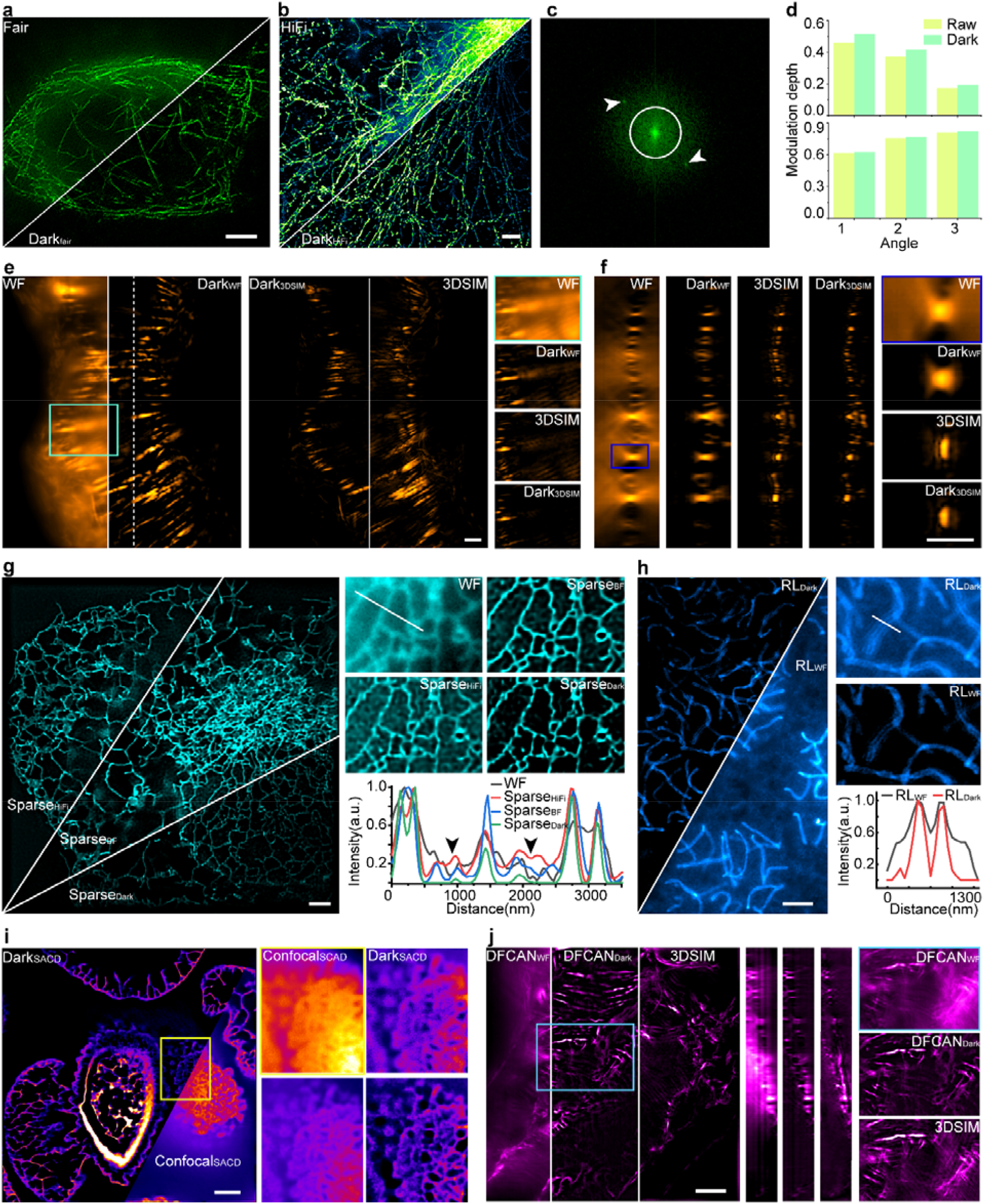
Dark sectioning assists various fluorescence image processing algorithms. The SIM result processed by Dark sectioning of (**a**) tubulin in U2OS (2D-SIM, fairSIM) and (**b**) tubulin in fixed Cos-7 cells (TRIF-SIM, HiFi-SIM) with (**c**) Fourier domain of 2D-SIM in (a). The white circle denotes the low-frequency part to be processed by Dark sectioning while the high-frequency part containing modulation points (white arrows) is not affected. (**d**) The modulation depth before and after Dark sectioning of three illumination angles.

### Dark Sectioning assists various fluorescence image processing algorithms

Furthermore, we demonstrate that Dark sectioning can be applied to different computational fluorescence imaging methods to improve their performance in case of severe background. We first did the comparison of 2D-SIM using the data from fairSIM in Fig. 4(a) and HiFi-SIM in Fig. 4(b). Even fairSIM and HiFi-SIM has widely recognized reconstruction performances, artifacts will appear when processing raw data with strong background. Applying Dark sectioning on the pre-processing of SIM’s raw data can reduce artifacts caused by the defocused background. The Fourier domain of fairSIM’s raw SIM data in Fig. 4(a) is shown in Fig. 4(c). Dark sectioning first divides the raw SIM image into the high-frequency part and low-frequency part bounded by a white circle in Fig. 5(c), and the high-frequency part is preserved without any change. Because the modulation points in the frequency domain lies in the high-frequency part, Dark sectioning will not change the pattern of SIM. This is also demonstrated by the modulation depth in Fig. 4(d), we can find that Dark sectioning will even slightly improve modulation depth because of the suppression of the low-frequency part of the SIM image. Then, we demonstrate that Dark sectioning can further improve the performance of 3DSIM using actin filament in mouse kidney section as shown in Fig. 4(e). Comparison between WF, Dark_WF_, 3DSIM, and 3DSIM processed by Dark sectioning (Dark_3DSIM_) shows that Dark sectioning can not only correctly distinguish and remove the defocused background of WF (proved by 3DSIM), but also greatly remove the artifacts of 3DSIM caused by the scattering background in the axial plane in Fig. 4(f). In addition, we demonstrate that Dark sectioning can help sparse deconvolution to eliminate artifacts caused by the background between adjacent ER tubes. Sparse deconvolution based on Dark sectioning (Sparse_Dark_) outperforms Sparse deconvolution based on HiFi-SIM(Sparse_HiFi_), Sparse deconvolution based on BF-SIM (Sparse_BF_) as shown in the black arrow which denotes the artifacts in Fig. 4(g). Furthermore, Dark sectioning can help Lucy-Richardson (RL) deconvolution to obtain a better resolution in the processing of the synaptonemal complex in Fig. 4(h) which may be submerged by the strong background. In addition, we apply Dark sectioning into SOFI using super-resolution imaging based on autocorrelation with a two-step deconvolution (SACD) algorithm^14^ as an example in Fig. 4(i). Although Dark sectioning cannot significantly improve the resolution on the basis of Confocal (resolution of Dark_confocal_ being 463.831nm, Confocal being 446.679 nm using rFRC^37^). Dark sectioning can strengthen the in-focus information to provide more information for SACD, so the resolution of SCAD when imaging thick samples like pine young staminate has been significantly improved from 353.314nm to 216.784nm using rFRC with a successfully SACD reconstruction (RSE of Dark_SACD_ being 0.0208, RSE of SACD being 0.0210) under the same parameters. Finally, we use the popular learning-based algorithm to process actin filament in the thick mouse kidney using DFCAN^38^ as an example in Fig. 4(j), finding DFCAN processed by Dark sectioning (DFCAN_Dark_) outperforms DFCAN based on WF images (DFCAN_WF_) in respect to background. Although DFCAN_Dark_ has a limited axial resolution, it has almost no artifacts compared with 3DSIM, thus providing a feasible method for volume image based on WF.

Upper denotes the modulation depth in (**a**) and bottom denotes the modulation depth in (**b**). (**e**) Dark sectioning can improve the performance of 3DSIM using actin filament in the mouse kidney as an example, with the comparison between WF, Dark_WF_, 3DSIM, and Dark_3DSIM_. (**f**) The axial plane of (**e**) denotes the fidelity of Dark sectioning and demonstrates the reduced artifacts of 3DSIM with its magnified zones. (**g**) Dark sectioning assists Sparse deconvolution when imaging ER tubes labeled with mCherry-Cytb5ER in living Cos-7 cells, and the corresponding comparisons between Sparse_HiFi_, Sparse_BF_, and Sparse_Dark_, with their profiles. (**f**) The axial plane of (**e**) denotes the fidelity of Dark sectioning and demonstrates the reduced artifacts of 3DSIM with its magnified zones. (**g**) Dark sectioning assists Sparse deconvolution when imaging near-core ER tubes, and the corresponding comparisons between Sparse_HiFi_, Sparse_BF_, and Sparse_Dark_, with their profiles. (**h**) Dark sectioning assists RL deconvolution to achieve better resolution of synaptonemal complex in mouse spermatoctyes, with its corresponding profile. (**i**) Dark sectioning helps super-resolution optical fluctuation imaging such as SCAD to achieve better resolution in case of high background using pine young staminate sample. (j) Dark sectioning helps learning-based methods such as DFCAN to approach 3DSIM with low artifacts using actin filament in mouse kidney section. Lateral scale bar: 4 μm. Axial scale: (**e**) 37 layers, 125 nm per layer. (**j**) 31 layers, 125 nm per layer.

## Discussion and conclusion

In this work, we have developed a versatile background removal algorithm called Dark sectioning that significantly improves optical sectioning in fluorescence microscopy. By exploiting dark channel priority and dual frequency separation, Dark sectioning reliably eliminates out-of-focus background while preserving weak in-focus signals across various microscopy modalities. Validation with confocal, 2p, and 3D-SIM shows that Dark Sectioning recovers fine details with high fidelity and greatly boosted signal-to-background ratio, structural similarity index measure, resolution-scaled error, and resolution-scaled Pearson coefficient compared to other methods.

Firstly, Dark sectioning improves optical sectioning across diverse fluorescence microscopy modalities including WF, SIM, STED, confocal, lightsheet, and light-field. It can enhance WF results close to confocal optical-sectioning results, 1p to 2p, and 2D-SIM to 3D-SIM. It recovers fine details even when in-focus signals are only 2% of overwhelming background. The output background-removal image provides 10 times higher SSIM and 10dB increased SBR. It outperforms existing computational methods like sliding paraboloid, rolling ball, sparse deconvolution, and commercial ICC, for background removal.

Secondly, it enables clearer observation of subcellular features by suppressing obscuring out-of-focus haze. In STED microscopy, scattering noise is effectively removed to reveal mitochondrial cristae ultrastructure. In lightsheet microscopy, mouse embryo can be visualized more clearly after Dark sectioning. For small-animal imaging of live mouse vascular, Dark sectioning can effectively remove the scattering background caused by mouse skin to enable sharp and accurate vascular mapping. In light-field microscopy imaging, Dark sectioning can help Vs-FLM better observe the 3D dynamic process of the neutrophils in mouse liver cells with almost no artifacts.

Finally, Dark Sectioning provides a simple, widely applicable solution to a fundamental fluorescence imaging limitation. It also complements other reconstruction/post-processing techniques. For 2D/3D-SIM, it improves illumination pattern contrast and reduce artifacts. For 3D-SIM, it further increases performance with fewer defocus artifacts. For polarized-SIM, it significantly improves polarization angle accuracy by removing backgrounds. Performance of other post-processing algorithms like sparse deconvolution, RL deconvolution, and machine learning methods like DFCAN, can all benefit from Dark Sectioning.

In some senses, the background removal algorithm is similar to the deconvolution algorithm. Deconvolution is to guess and add high-frequency parts beyond the diffraction limit, while background removal is to subtract and minus low-frequency parts as background. However, background removal algorithms have been long dragged by deconvolution algorithms such as RL, Sparse, or MRA deconvolution where statistics, sparse, continuity, or wavelet priority are introduced. Background removal algorithms are still based on the assumption that background lies in the low-frequency parts in the Fourier or wavelet domain. Here, we introduce dark channel priority into fluorescence images, then combine it with dual frequency separation and physical characteristics of PSF. The high fidelity of Dark sectioning is proved by various methods such as 3DSIM, 2p, and confocal. Comparing with the deep learning approaches in fluorescence microscopies which are often “black-box” and heavily training dependent and parameter empirical tunning, Dark sectioning has a transparent and strict mathematical process, providing high-fidelity results for new biological discoveries.

Dark sectioning provides a powerful solution to improve optical sectioning capability that is compatible with most fluorescence microscopic modalities, through overcoming a major limitation of defocused background. We provide easily accessible, open-source packages with MATLAB, Fiji, and Exe versions to facilitate the community. We emphasize that Dark sectioning is a common method for different microscopies and various algorithms. Our work may break the tradeoff between defocus background and lower temporal resolution with complex setup, thus enabling clearer visualization of subcellular features for biological discovery.

## Acknowledgments

P. X. acknowledges the funding support from the National Key R&D Program of China (2022YFC3401100) and the National Natural Science Foundation of China (62025501, 92150301). We thank the National Center for Protein Sciences and the Core Facilities of Life Sciences at Peking University. We thank Ziang Chen in Shandong University for the culture of mouse spermatocytes and label of synaptonemal complex.

## Author Contributions Statement

P. X. and J. Q. supervised the project. R. C., Y. L., J. Q., and P. X. initiated and conceived the research. R. C. developed the algorithm and implemented the software. Y. L. construct the multi-mode SIM system. W. W. conduct the live cell imaging experiment. G. Z. and J. W. conduct the one/two-photon joint experiment. G. W. conduct the THUNDER experiment. Y. S. and C. L. conduct the small animal imaging experiment. W. R. conduct the spinning disk experiment. W. R. conduct the spinning disk system and experiment. J. S. conduct the STED imaging experiment. Y. H. helps the reconstruction of deepMRA. X. X. and M. L. give the analysis and discussion of data. J. H. and Y. L. conduct the DFCAN reconstruction. R. C. and Y. L. draw the figures and videos. R. C., Y. L., J. Q., and P. X. write the manuscript with the input from all authors.

## Code availability

Software of Dark sectioning is provided to reviewers and uploaded to Github: https://github.com/Cao-ruijie/Dark-sectioning. This software contains MATLAB, Fiji, and Exe version with detailed user guide for users.

## Data availability

Typical data in this article is provided to reviewers and uploaded to Figshare: https://figshare.com/articles/dataset/Dark-sectioning/24607614. This data contains detailed parameters and comparisons.

## Methods

### Commercial samples

The mouse tails sections in Fig. 1(d) are purchased from Voyage of Discovery Teaching Equipment (China). The samples of aspergillus conidiophores in Fig. 1(e) is purchased from Carolina Biological Supply (297872, American). The mouse kidney sections in Fig. 3(a), Fig. 4(e, f, j) are purchased from ThermoFisher (F24630, American). Pine young staminate sample in Fig. 4(i) is purchased from Suzhou Shenying Optical co Itd.

### Open-source data

The mouse kidney sections in Fig. 3(a), Fig. 4(j) are from our previously proposed DMD-3DSIM^30^. Lightsheet data Fig. 3(e, f) are from practical considerations for quantitative light sheet fluorescence microscopy^35^ https://doi.org/10.6084/m9.figshare.c.6211429. The neutrophils with high-speed migration in living mouse in Fig. 3(g, h) are from the Bio-LFSR^36^ dataset https://doi.org/10.5281/zenodo.7233421. Tubulin in U2OS cells in Fig. 4(a) is from fairSIM^17^: https://www.fairsim.org/. Tubulin in COS-7 cells in Fig. 4(b) is from HiFi-SIM^16^: https://www.nature.com/articles/s41377-021-00513-w.

### Open-source algorithms

In Fig. 1(d), we use the dark priority in haze removal (for natural outdoor images) algorithm^26^ in https://github.com/sjtrny/Dark-Channel-Haze-Removal. In Fig. 2(b), Fig. 3(a), Fig. 4(e), Open-3DSIM^11^ (v2.2) is used for 3DSIM reconstruction in https://github.com/Cao-ruijie/Open3DSIM. In Fig. 2(b), Fig. 4(b), Fig. 4(g), HiFi-SIM^16^ (v1.01) is used in https://www.nature.com/articles/s41377-021-00513-w. In Fig. 2(i), Weka segmentation^33^ is trained and used by trainable Weka segmentation plugin in Fiji. In Fig. 4(g), BF-SIM^5^ (v1.2) is used in https://doi.org/10.6084/m9.figshare.22640974.v1. In Fig. 4(g), VsLFM^36^ (v3.1) is used in https://github.com/THU-IBCS/VsLFM-master. In Fig. 5(b), fairSIM^17^ is used in https://www.fairsim.org/. In Fig. 4(i), SCAD^14^ (v0.2.0) with 100 frames raw images is used in https://github.com/WeisongZhao/SACDm. In Fig. 4(j), DFCAN^38^ is used in https://github.com/qc17-THU/DL-SR.

### Cells culture

We seed COS-7 cells in 8 well chamber sliders, wait until the cells grow to 70%-90% confluence. To label mitochondria, we use complete cell culture medium to mix PKmito RED (Cytoskeleton, CY-SC052) was diluted to 200 times, and replace the original medium in the chamber, and then the cells were incubated at 37°C in a 5% carbon dioxide incubator for 15 minutes for SIM imaging in 640 nm channel. And SiRPFA was diluted to 500nM and used to label mitochondria before drug-induced apoptosis and Rapamycin was used to induce apoptosis after 20 minutes for STED imaging in 637 nm channel. To label tubulin and lysosomal, we first use SiR Tubulin Kit^39^ (Cytoskeleton, CY-SC002) to label tubulin in live COS-7 cells under the concentration of 1μM for 1 hour at 37°C. We dilute LysoView™ 488(Biotium,70067) for 1000 times to label lysosomal in live COS-7 cells for 30 minutes at 37°C. Tubulin and lysosomal are imaged in 640 nm and 488 nm channels. To transfer ER tubes, we take 5 μL of Opti-MEM culture media (Gibco, 31985070) in a centrifuge tube, add 0.5 μL of Lipofectamine 3000 (Invitrogen, L3000150) and mix well as working solution A. Take 5 μL Opti-MEM culture media, 0.2 μL P3000 reagent, 1 μL 100ng/μL GFP-KDEL plasmid DNA, mix well and use it as working solution B. Mix working solution A and working solution B, and let stand at room temperature for 10-15min. Add 10 μL of the mixed solution to each well into 8 Well Chamber sliders, and use a pipette to mix well, incubate at 37°C, 5% carbon dioxide in a cell culture incubator for 6h-8h, then remove the medium containing DNA, Replace with the complete medium for culturing cells, take it out after continuing to incubate for 12-24 hours, and use channel 488 for imaging. The protocol to culture mouse spermatocytes and label synaptonemal complex are accomplished by Shandong University, following ref^40^. The culture of human tonsil tissues and label of nuclei is accomplished by Alpha X (Beijing) Biotech CO., LTD, China. Human tonsil tissues were incubated for 1 h at 37°C. Then slides were incubated for 10 min at 37°C.

### DMD-based multi-mode joint SIM microscope

To validate Dark sectioning is reliable and can optimize various SIM-based methods, we construct a DMD-based multi-modality joint SIM system containing the modalities of WF, TIRF, HiLo, 2DSIM, and 3DSIM. The excitation laser beam of 561 nm was polarized via a Glan-Taylor polarizing prism and modulated by a polarization rotator composed of an electro-optic modulator and a quarter wave plate. The emergent beam was collimated and expanded via a beam expander and then illuminated on the pattern generator Digital micro-mirror device (DMD). The diffraction beams converging through lens1 form multiple diffraction orders. The custom-built spatial mask is located in a pupil plane of lens1 and can filter the needful order diffraction beams (the 0^th^ and ±1^st^ orders in 3DSIM, the +1^st^ and -1^st^ orders in 2DSIM, the +1^st^ and -1^st^ orders in HiLo, the +1^st^ or -1^st^ order in TIRF, and the 0^th^ order in WF). The diffraction beams were refocused to back focal plane (the center and two points near the opposite edges in 3DSIM, two points near the opposite edges in 2DSIM, two points with the opposite direction in HiLo, one point near the edge in TIRF, and the center point in WF) of the objective lens through a 4f system consisting of lens2 and lens3 via a polarization-preserving dichroic mirror. The objective recollimated the three beams and made them approach the sample and intersect with each other to produce a structured pattern of excitation intensity in the focal plane of the objective. The fluorescence emitted from the sample was gathered by the same objective and passed through the same polarization-preserving dichroic mirror, a tube lens, and an emission filter to a water-cooled sCMOS camera to detect the emission fluorescence.

### One-photon and two-photon joint validation

To validate Dark sectioning can remove backgrounds with high fidelity and correctly subtract neuronal activities, we use our previously built joint one-photon and two-photon detection system based on standard TPLSM, and a 488-nm-centered wide-field illumination path and a camera detection path are added to the system. The details can be seen in DeepWonder.

### WF and confocal joint validation

To validate Dark sectioning is commonly useful for WF images, we use a Dragonfly microscopy (Andor Technology, England) for wide-field and confocal joint imaging. Dragonfly is a spinning disk system for confocal imaging equipped with 405/488/561/637 nm lasers, 100X/1.44 objective, and sCMOS from Zyla.

### STED imaging

We use STED super-resolution confocal microscope (TCS SP8 STED 3X, Germany) for imaging. The laser wavelength used was 637 nm, and STED depletion was achieved with a pulsed laser at 775 nm. A 100× oil-immersion objective lens was used for imaging.

### SOFI imaging

We use our home-built spinning disk confocal microscopy to conduct SOFI. A 4-independently-controllable integrated light source was used (CELESTA Light Engine, Lumencor) as the illumination laser. Our imaging configuration included a 100× objective lens (CFI SR HP Apo TIRF 100×, 1.49NA, Nikon) in conjunction with a sCMOS camera (ORCA-Flash4.0 V3, Hamamatsu). The spinning disk maintains a consistent speed of 5000 rpm.

### WF and 2DSIM imaging

We use our custom-built commercial Ariy-SIM system for WF and 2DSIM live cells imaging. Ariy-SIM is equipped with 405nm, 488nm, 561nm, 640nm channels with an oil objective from Nikon.

### Thunder imaging

We conduct experiments for comparison between Dark sectioning and Leica’s ICC algorithm using THUNDER Imager Tissue. THUNDER Imager Tissue is a wide-field microscopy with ICC algorithm to remove background at 561/640 nm excitation wavelength with an objective of 0.95 NA.

### Small-animal imaging

We selected indocyanine green (ICG), which has been used in clinical practice, as a fluorescent dye and injected into mice through tail vein injection. A 785 nm continuous laser (CNI MDL-III-785, China) was used to uniformly irradiate the mouse body through a 1/2 fiber bundle to stimulate fluorescence signals. We use a cooled infrared fluorescence camera (Hamamatsu C11440, Japan) to collect fluorescence signals and place a filter in front of the camera lens to filter out background light.

### Signal-to-background ratio

We use the signal-to-background ratio (SBR) to evaluate the background suppression capability of different algorithms. First, we normalize the signal distribution from the original image *I x, y* to the normalized image *I* ‘(*x, y*):

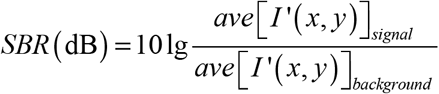

Where *ave*[·] is the average gray value of the region of interest of signal or background. For fair comparisons, we choose the same area of signal or background using different algorithms.

### Signal-to-noise ratio

We use the signal-to-noise ratio (SNR) to evaluate the ability to remove the defocused background of different algorithms. We choose the part with signal and the part without signal to solve its mean value and variance, and use the following formula to solve SNR (dB):

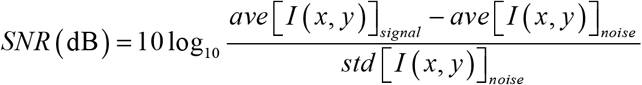

Where *ave* [·] and *std* [·] are the average gray value and standard deviation of the region of interest. For fair comparisons, we choose the same area of signal or noise using different algorithms.

### Structure similarity index measure

We use the structure similarity index measure (SSIM) to evaluate the similarity between the image *I* (*x, y*)and GT *G* (*x, y*) image. The SSIM between two images can be expressed as:

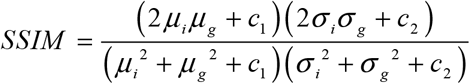

Where *μ*_*i*_ and *μ*_*g*_ are the average value of *I* (*x, y*) and *G* (*x, y*). *σ*_*i*_and *σ*_*g*_ are their standard deviation value. *c*_1_ and *c*_2_ are the constants which can be expressed as *c*_1_ *=*(*k*_1_ *L*)^2^ and *c*_2_ *=*(*k*_2_ *L*)^2^, where *L* is the pixel number of the image, and *k*_1_ 0.01, *k*_2_ 0.03. When SSIM is close to 1, it indicates that the deviation between two images is small.

### Resolution-scaled error and Resolution-scaled pearson coefficient

NanoJ-SQUIRREL^29^ is used to evaluate the bias between the GT image and the reconstructed image. Resolution-scaled error (RSE) represents the sum of error between the scaled GT image and reconstructed image, and resolution-scaled pearson coefficient (RSP) represents the pearson coefficient between the scaled GT image and reconstructed image. The closer the RSP is to 1, the closer the RSE is to 0, indicating that the GT and reconstructed images are closer.

### Correlation score

We use Pearson’s correlation coefficient to evaluate the similarity of temporal activities between one-photon and one-photon with Dark sectioning using two-photon microscopy as GT. When Pearson’s correlation coefficient is closer to 1, it means the image is closer to the GT.

### rFRC mapping

We use rFRC mapping^37^ to analysis the resolution of SOFI/SACD. The rFRC map denotes the spatial distribution of resolution. And the RSE denotes the difference between low-resolution and high-resolution images. The Dark_SACD_ and SACD images are using two high-resolution and one low-resolution images to calculate. Confocal and Dark_Confocal_ images are using single frame resolution calculation with 1/7 threshold.

### Imaging process and analysis

We use Fiji to conduct MIP in *xoy* plane, intensity histogram, Weka segmentation, SBR calculation, SNR calculation, NanoJ-SQUIRREL, rolling ball, and sliding paraboloid algorithms. We use MATLAB to conduct the SSIM, Pearson’s correlation coefficient, and MIP in xoz plane. Origin is used to plot curve, Adobe Illustrator is used to draw the figures, PowerPoint are used to make videos, and the brightness and contrast are adjusted linearly for display purpose.

